## Supplementary figures and images for "The Hippo kinase cascade regulates a contractile cell behavior and cell density in a close unicellular relative of animals"

### Figure 1-supplement 1-source data 2.tif

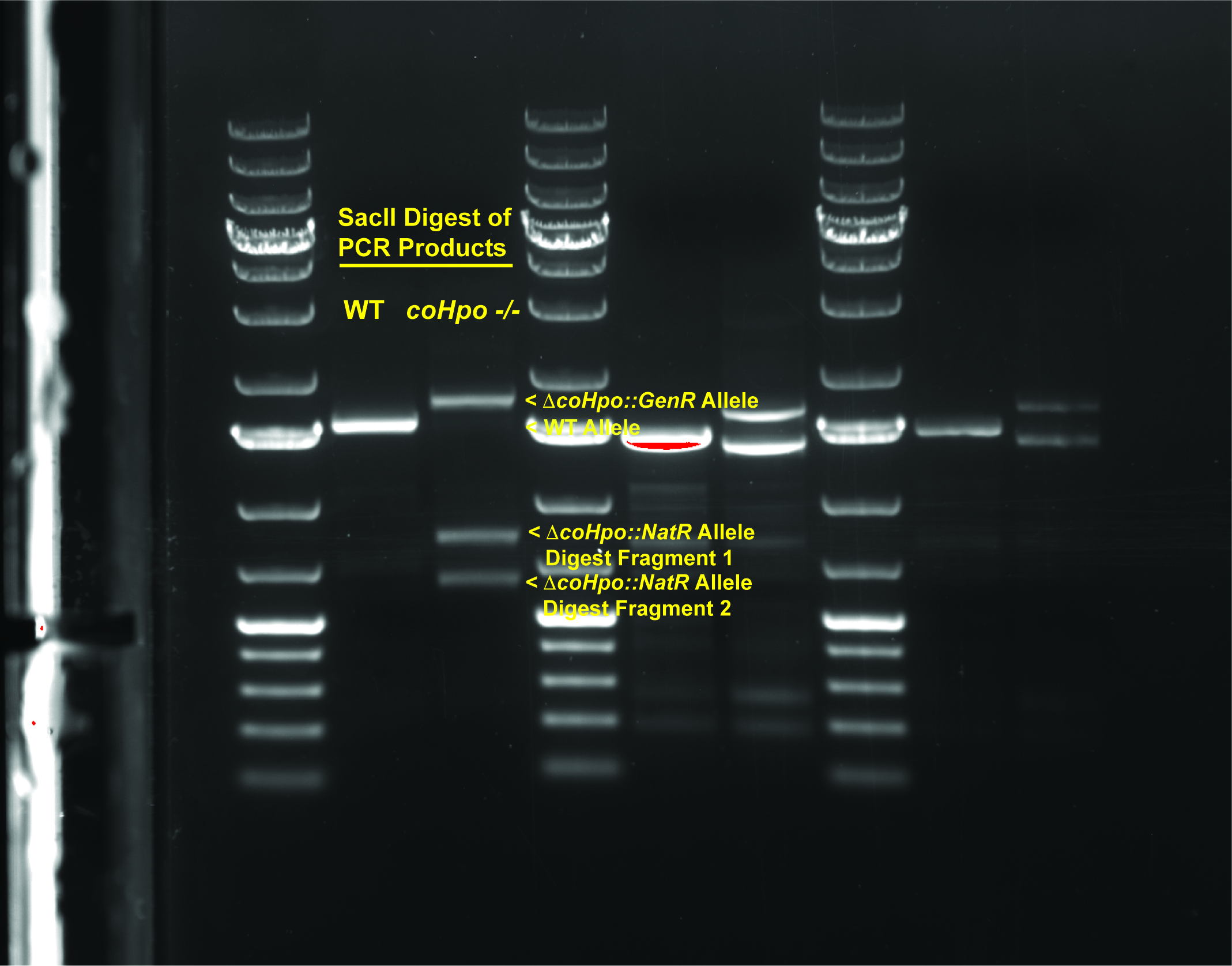

### Figure 1-supplement 1-source data 3.tif

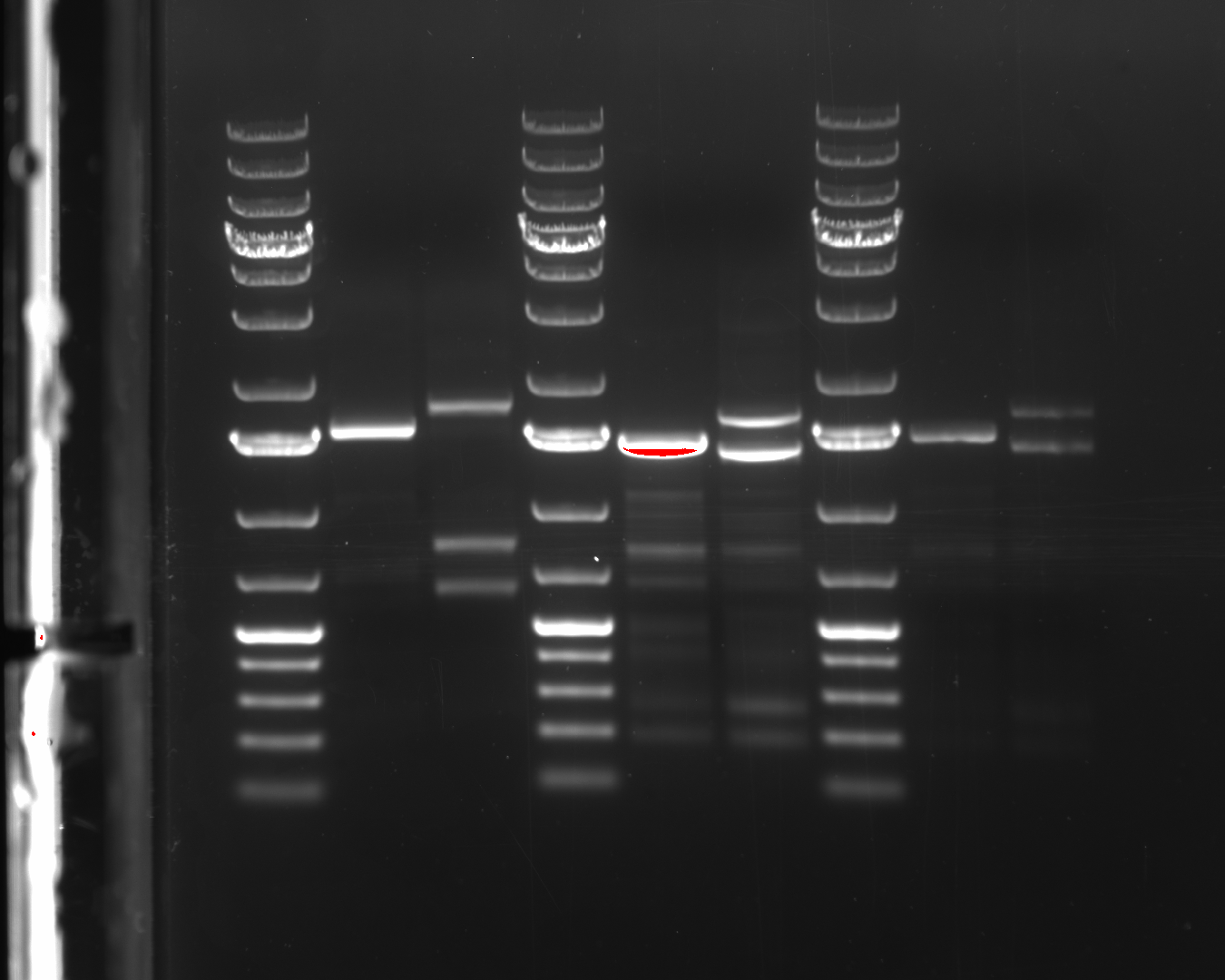

### Figure 1-supplement 1-source data 4.tif

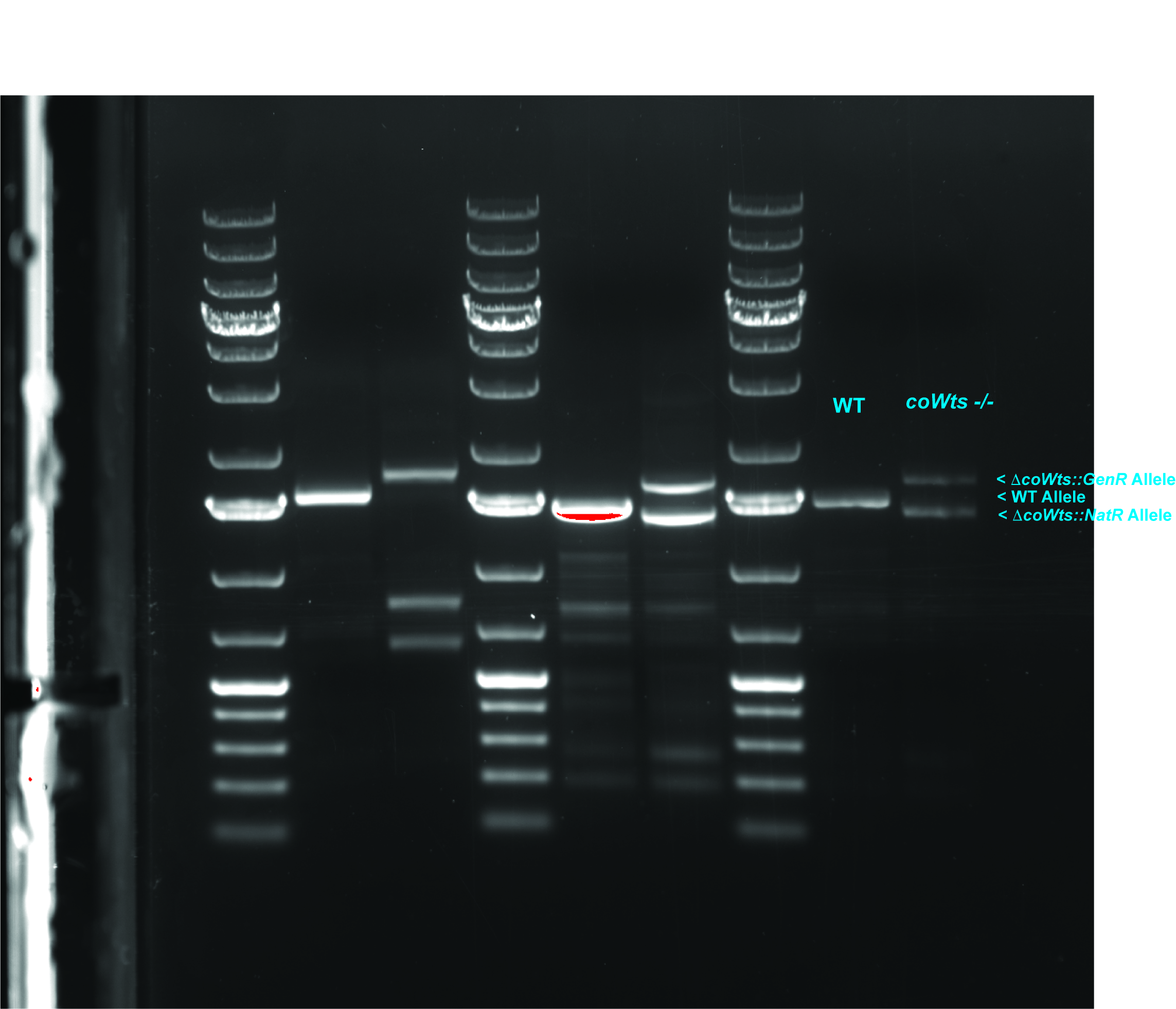
